# Impacts of the feedback loop between sense-antisense RNAs in regulating circadian rhythms

**DOI:** 10.1101/2024.04.28.591560

**Authors:** Koichiro Uriu, Juan P. Hernandez-Sanchez, Shihoko Kojima

**Affiliations:** School of Life Science and Technology, Tokyo Institute of Technology, Meguro, Tokyo, Japan; Graduate School of Natural Science and Technology, Kanazawa University, Kanazawa, Ishikawa, Japan; Department of Biological Sciences, Fralin Life Sciences Institute, Virginia Tech, Blacksburg, VA, USA

**Keywords:** circadian rhythms, antisense RNA, transcriptional interference, amplitude, robustness, mathematical model

## Abstract

Antisense transcripts are a unique group of non-coding RNAs that are transcribed from the opposite strand of a sense coding gene in an antisense orientation. Even though they do not encode a protein, these transcripts play a regulatory role in a variety of biological processes, including circadian rhythms. We and others found an antisense transcript, *Per2AS*, that is transcribed from the strand opposite the sense transcript *Period2* (*Per2*) and exhibits a rhythmic and antiphasic expression pattern compared to *Per2* in mouse. By assuming that *Per2AS* and *Per2* mutually repress each other, our previous mathematical model predicted that *Per2AS* regulates the robustness and the amplitude of circadian rhythms. In this study, we revised our previous model and developed a new mathematical model that mechanistically described the mutually repressive relationship between *Per2* and *Per2AS* via transcriptional interference. We found that the simulation results are largely consistent with experimental observations including the counterintuitive ones that could not be fully explained by our previous model. These results indicate that our revised model serves as a foundation to build more detailed models in the future to better understand the impact of *Per2AS-Per2* interaction in the mammalian circadian clock. Our mechanistic description of *Per2AS-Per2* interaction can also be extended to other mathematical models that involve sense-antisense RNA pairs that mutually repress each other.

## Introduction

Transcription is pervasive in mammalian genome, and up to 80% of the genome is actively transcribed even though protein-coding genes account for only about 2% (*Djebali et al., 2012, Nurk et al., 2022*). Many of these transcripts do not encode a protein but can exhibit a wide spectrum of functions in diverse biological and pathophysiological processes such as cell cycle and cancer, X inactivation, imprinting, neurological pathways, and immune responses (*Beiter et al., 2009, Mosig and Kojima, 2022, Wanowska et al., 2018*). Currently, three major mechanistic possibilities are proposed to describe how these transcripts function without encoding a protein. First is the transcript model, in which transcripts themselves are the functional biomolecule and interact with DNA/chromatin, other RNA, or protein to exhibit functions (*Ali and Grote, 2020, Mosig and Kojima, 2022*). Second is the DNA model, in which a gene regulatory element is embedded in the DNA of the non-coding transcript’s locus and the transcriptional activity of the non-coding RNA directs the activity of the DNA regulatory element, ultimately regulating the expression of target genes. Third is the transcription model, in which the process of transcription of a non-coding RNA, rather than the RNA product, regulates the transcriptional activity of a neighboring genes (*Ali and Grote, 2020, Mosig and Kojima, 2022*).

These non-coding transcripts can be further categorized into different types by their genomic location and orientation in relation to nearby protein coding genes (*Mosig and Kojima, 2022*). Of those, we are particularly interested in antisense transcripts that are transcribed from the opposite strand of a coding gene locus in an antisense orientation (*Bertone et al., 2004, Carninci et al., 2005, Katayama et al., 2005, Røsok and Sioud, 2004, Sun et al., 2006*). In mammals, 25-40% of protein-coding genes have antisense transcript partners and their expression pattern can be concordant or discordant with that of their sense partner RNAs (*Katayama et al., 2005, Sun et al., 2006*). Antisense transcripts of the sense coding genes that are critical for circadian rhythms have been reported in *Neurospora crassa* and *Antheraea pernyi* (*Kramer et al., 2003, Li et al., 2015, Mosig et al., 2021, Sauman and Reppert, 1996, Xue et al., 2014*), more recently in *Mus musculus* (*Koike et al., 2012, Menet et al., 2012, Vollmers et al., 2012, Zhang et al., 2014*). Interestingly, the expression of sense core clock mRNAs and their antisense transcript partners are all rhythmic and antiphasic (*Koike et al., 2012, Kramer et al., 2003, Menet et al., 2012, Sauman and Reppert, 1996, Vollmers et al., 2012, Zhang et al., 2014*), leading to the hypothesis that sense and antisense RNAs mutually repress each other.

We previously constructed mathematical models to understand the function of an antisense transcript, *Per2AS*, to the sense *Period2* (*Per2*) coding gene in the mammalian circadian circuit (*Battogtokh et al., 2018, Mosig et al., 2021*). As a core circadian clock gene, *Per2* negatively regulates its own transcription, contributing to the generation of self-sustained circadian oscillation (*Takahashi, 2017, Zheng et al., 2001*). When we assumed that *Per2* and *Per2AS* repress each other, the model predicted that the mutual repression between *Per2AS* and *Per2* would be critical in conferring robustness to the circadian system (*Battogtokh et al., 2018, Mosig et al., 2021*). This was further corroborated with an experimental observation, in which *Per2AS* regulates the amplitude of the circadian clock in mouse fibroblasts (*Mosig et al., 2021*). At the same time, our model did not seem to fully explain other experimental observations.

In this study, we revised our previous model and described the regulatory relationship between *Per2AS* and *Per2* more mechanistically. We found that our new model can explain many experimental observations, including counterintuitive ones, supporting the idea that the mutual repression between *Per2AS* and *Per2* is a feasible mechanism. However, we also found additional experimental conditions that cannot be easily explained by the revised model, suggesting the need for further refinement.

## Results

### Mechanistic description of regulatory relationships between Per2AS and Per2

In our previous work, we developed two mathematical models to understand the functions of *Per2AS*. The first model is more comprehensive and includes many core clock genes and feedback loops, whereas the second model is much simpler and only includes *Per2AS* and *Per2* (*Battogtokh et al., 2018, Mosig et al., 2021*). In both models, we assumed that *Per2AS* and *Per2* mutually repress each other and the repression is mediated by their RNA products. However, our experimental observations suggested that the repression occurs independent of their RNA and/or protein products and in fact the act of transcription on one strand suppresses the transcription of the other strand, a mechanism called transcriptional interference (*Georg and Hess, 2011, Mosig et al., 2021, Shearwin et al., 2005, Xue et al., 2014*).

To improve our previous models, we decided to use a simpler version of our model that only includes *Per2AS, Per2*, and PER2, as a starting point (Fig. 1A) (*Mosig et al., 2021*). We then focused on describing the regulatory relationship between *Per2AS* and *Per2* more mechanistically. To this end, we first assigned variables that describe the state of transcription (*Sneppen et al., 2005*). We defined variables *X*_*S*_(*t*) and *X*_*A*_(*t*) (0 ≤ *X*_*i*_ ≤ 1, *i* ∈ {*S, A*}) as transcriptional activities of *Per2* and *Per2AS*, respectively. They represent a condition in which RNA polymerase (RNAP) is recruited to the promoter region at time *t* (Fig. 1B: left, ‘ON’) and RNAPs start transcription immediately after its recruitment. Accordingly, the state of inactive RNA transcription (i.e., RNAP not being recruited to the promoters) at time *t* can be described as 1 − *X*_*i*_(*t*) (Fig. 1B: right, ‘OFF’).

**Figure 1.**
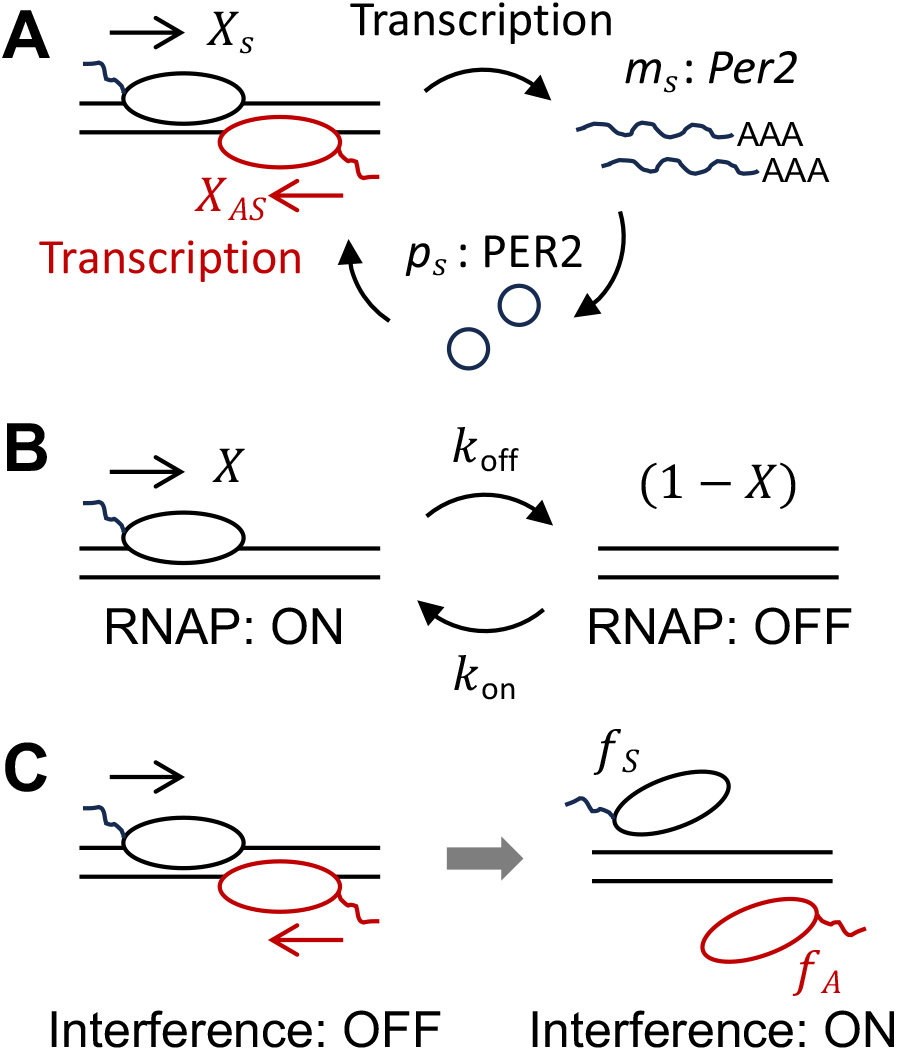
Mechanistic description of the regulatory relationship between *Per2AS* and *Per2*. (A) Autoinhibitory feedback loop of *Per2. Per2* mRNA (*m*_*S*_) is transcribed from DNA with transcriptional activity *X*_*S*_. It is ultimately translated into PER2 protein (*p*_*S*_), which inhibits its own transcription. *Per2AS* is transcribed with its activity *X*_*A*_. Oval represents RNA polymerase (RNAP). (B) Description of the RNAP recruitment and transcriptional activity. RNAP is recruited to DNA (left: ON) with the rate *k*_on_, during which transcription is always active. RNAP is detached from DNA with the rate *k*_off_, during which no RNA is transcribed (right: OFF). (C) Description of transcriptional interference by RNAP collision. When two RNAPs on opposite strands are both recruited to DNA and transcribing RNAs in the opposite direction, they collide each other with probability *X*_*S*_ & *X*_*A*_. This results in both RNAPs being detached from DNA with probabilities *f*_*S*_ and *f*_*A*_, and no RNAs can be transcribed.

We also described the transitions of RNAP recruitment between ‘ON’ and ‘OFF’ (Fig. 1B) at sense and antisense strands as:

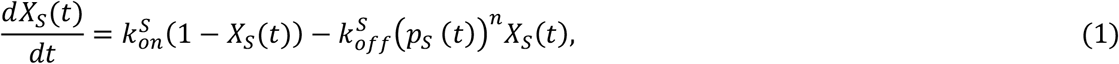

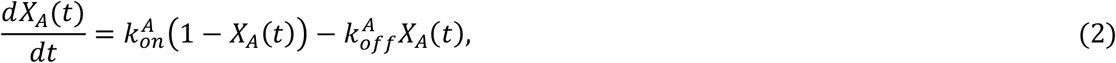

where *p*_*S*_ (*t*) is the PER2 protein level at time *t*. Note that this is a dimensionless variable (see Methods), while we preserved the timescale unit as hour because this is the characteristic timescale of the circadian clock system (i.e., 24 hours). The first terms in Eqs. (1) and (2) represent the dynamics of RNAP recruitment to DNA with the rate 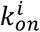(Fig. 1B: from OFF to ON), while the second terms represent RNAP detachment from DNA at the rate 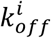 (Fig. 1B: from ON to OFF). The transition from ON to OFF for *Per2* depends on the PER2 protein level with nonlinearity *n*, because of its autoinhibition (Fig. 1A). In contrast, the time derivative of *X*_*A*_ Eq. (2) is independent of both *Per2* mRNA and PER2 protein levels.

We next considered the mutual repression between *Per2AS* and *Per2* by transcriptional interference, in which RNAPs on the opposite strands and opposite directions collide (Fig. 1C). To represent this condition, we described the dynamics of *Per2* mRNA *m*_*S*_ and *Per2AS m*_*A*_ as:

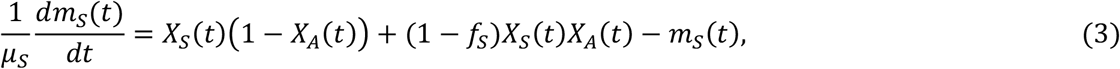

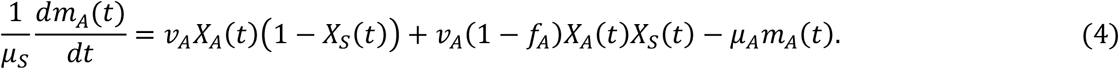

Note that *m*_*S*_, *m*_*A*_ and *p*_*S*_ are all dimensionless variables (see Methods for derivation). Eq. (3) describes the dynamics of *Per2* mRNA, including *Per2* mRNA synthesis when *Per2AS* is not transcribed (the first term) or transcribed (the second term) and its degradation (the third term). Eq. (4) represents the dynamics of *Per2AS* and each term corresponds to Eq. (3), although on the antisense strand. We also introduced the probability of transcriptional interference as *f*_*S*_ or *f*_*A*_, in which RNAPs collide and get detached from sense or antisense DNA, respectively (Fig. 1C). If *f*_*A*_ = 0, there is no RNAP detachment from the *Per2AS* strand by collision. In contrast, if *f*_*A*_ = 1, RNAPs are always detached by RNAP collision when RNAP is recruited to DNA. We assumed that RNAP collision would result in incomplete transcription (*Bordoy and Chatterjee, 2015, Sneppen et al., 2005*).

We also described the dynamics of PER2 protein level as:

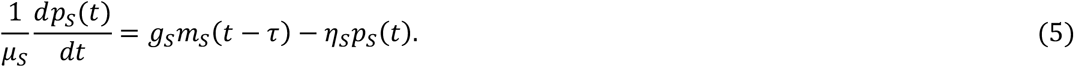

The first term represents the production of functional PER2 protein at the translation rate *g*_*S*_ from a full-length *Per2* mRNA, whereas the second term represents the degradation of PER2 protein at a rate *η*_*S*_ (see Methods). We also used the parameter *τ* to collectively describe the time delay required for PER2 to ultimately inhibit its own transcription (*Korenčič et al., 2012, Uriu and Tei, 2021*).

### Quasi-steady state analysis reveals that PER2 protein promotes Per2AS transcription

We next performed quasi-steady state analysis to simplify the equations and reduce the number of independent variables, given that the timescale of RNAP recruitment and detachment from DNA is much shorter than that of the circadian clock (*Lammers et al., 2020, Meeussen and Lenstra, 2024*). By setting *dX*_*S*_/*dt* = *dX*_*A*_/*dt* = 0 in Eqs. (1) and (2), we obtain:

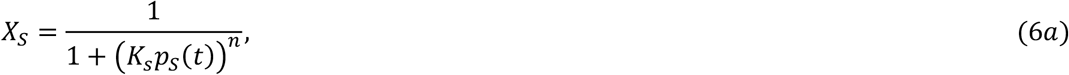

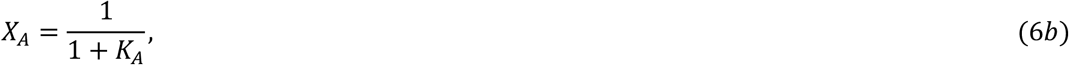

Where 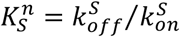 and 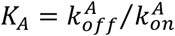 are the ratios of the transition rate between RNAP detachment and recruitment on the *Per2* and *Per2AS* strands, respectively. By using Eq. (6) in Eqs. (3) and (4), we obtain the dynamics of *Per2* and *Per2AS* level as:

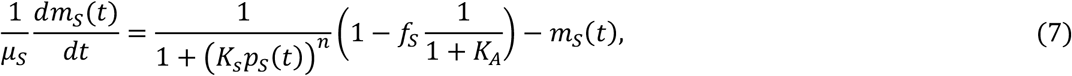

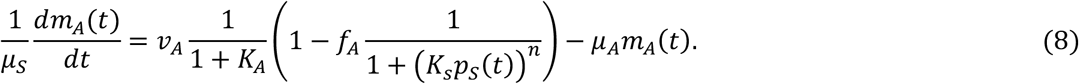

Eq. (7) shows that the value of *K*_*A*_ affects the transcription of *Per2* mRNA depending on *f*_*S*_. Eq. (8) reveals that the PER2 protein level promotes the transcription of *Per2AS*, even though the original equations for *Per2AS* dynamics (Eqs. (2) and (4)) do not include any variables deriving from PER2.

### Our new model with transcriptional interference successfully describes the antiphasic expression patterns of Per2 and Per2AS

Eq. (8) suggests that if PER2 protein level *p*_*S*_ is rhythmic, it can drive the rhythmicity of *Per2AS*. In addition, PER2 can also drive antiphasic patterns between *Per2AS* and *Per2*, because while PER2 protein represses *Per2* mRNA (Eq. (7)), it activates *Per2AS* (Eq. (8)). To test this, we numerically solved the delay differential equations (DDEs) (5), (7) and (8) (Methods) using parameter values based on the experimental observation whenever possible (Table 1). For example, we set the translation rate of PER2 from a single mRNA as 14 per hour, given the average speed of translation (5 amino acids per second) and the length of PER2 protein (1257 aa) (*Wu et al., 2016*). We also set *τ* = 6.2 h given that there is a 4 - 6 hour phase lag between the peaks of *Per2* mRNA and PER2 protein (*Field et al., 2000*). When no information is available, we set values similar to the corresponding parameters that are already known. For example, we used the same transcription rate for both *Per2AS* and *Per2* (*v*_*A*_ = 1) but determined the value of *K*_*A*_ so that the peak *Per2AS* level is approximately a third of the peak *Per2* level as was observed in mouse liver (*Koike et al., 2012*). We also estimated the half-life of *Per2AS* as 2 hours, the same as that of *Per2* and PER2 (*Friedel et al., 2009, Matsumura et al., 2022, Sharova et al., 2009*). We also considered the simplest and most plausible situation in which RNAPs on both strands detach from DNA by collision with the same probability *f*_*S*_ = *f*_*A*_ = *f*.

**Table 1.** List of parameters used in the mathematical model

| parameter | value* | description | References |
| --- | --- | --- | --- |
| $n$ | 3 | exponent of PER2 protein in on-to-off transition of *Per2* transcriptional activity | - |
| $f_S$ | - | Probability of RNAP collision and detachment for *Per2* | - |
| $\mu_S$ | $\log(2)/2$ | degradation rate of *Per2* mRNA | (Friedel et al. , 2009, Sharova et al. , 2009) |
| $K_S$ | 0.3684 | $n$th root of the ratio of RNAP detachment rate to recruitment rate for *Per2* | - |
| $v_A$ | 1 | relative transcription rate of *Per2AS* | - |
| $f_A$ | - | Probability of RNAP collision and detachment for *Per2AS* | - |
| $\mu_A$ | 1 | relative degradation rate of *Per2AS* | - |
| $K_A$ | 4.0 | ratio of RNAP detachment rate to recruitment rate for *Per2AS* | - |
| $g_S$ | $14/\mu_S$ | relative translation rate of PER2 protein | (Wu et al. , 2016) |
| $\eta_S$ | 1 | relative degradation rate of PER2 protein | (Matsumura et al. , 2022) |
| $\tau$ | 6.2 | time delay of functional PER2 protein production | (Field et al. , 2000) |

\*: used in figures for normal condition. -: specified in each figures.

When there is no interference (*f* = 0), the level of *Per2AS* converges to a steady state *v*_*A*_/(*μ*_*A*_(1 + *K*_*A*_)), while *Per2* mRNA and PER2 protein oscillate with a period of 23.87 hrs because of its autoinhibitory feedback loop (Fig. 2A). The autonomous *Per2* and PER2 rhythms without interference by *Per2AS* is consistent with the observations that some tissues maintain circadian rhythms in the absence of *Per2AS* (*Mosig et al., 2021*). When the probability of RNAP collision and detachment is *f* > 0, the expression of *Per2AS* becomes rhythmic (Fig. 2B, C). When the limit cycle trajectory is projected onto the *Per2* mRNA-*Per2AS* concentration space with *f* = 1 to understand their phase relationship, we found that the expression patterns of *Per2AS* and *Per2* are antiphasic as was reported previously (*Koike et al., 2012, Menet et al., 2012, Vollmers et al., 2012, Zhang et al., 2014*), because *Per2AS* level becomes highest when *Per2* mRNA level is the lowest and vice versa (Fig. 2D). The trajectory on the PER2-*Per2AS* concentration space (Fig. 2E), on the other hand, demonstrates that *Per2AS* level decreases and increases together with PER2 protein level (left downward and right upward arrows in Fig. 2E), and becomes highest after the PER2 protein peak (left upward arrow). As the probability of RNAP collision and detachment *f* increases from 0 to 1, the lowest level of *Per2AS* (trough of oscillation) decreases (Fig. 2F: red dotted line) while its highest level (peak of oscillation) remains unchanged (Fig. 2F: red solid line), resulting in the increase of the amplitude of *Per2AS*. In contrast, the highest level of *Per2* mRNA decreases while the lowest level remains almost the same as *f* increases, resulting in a reduction of amplitude of the *Per2* mRNA oscillation (Fig. 2F: bottom). We also found that the period of *Per2* mRNA became shorter as *f* increased (Fig. 2G). The period of *Per2AS* is the same as that of *Per2* because the oscillation of *Per2AS* level is driven by *Per2* via transcriptional interference (Fig. 2A-C).

**Figure 2:**
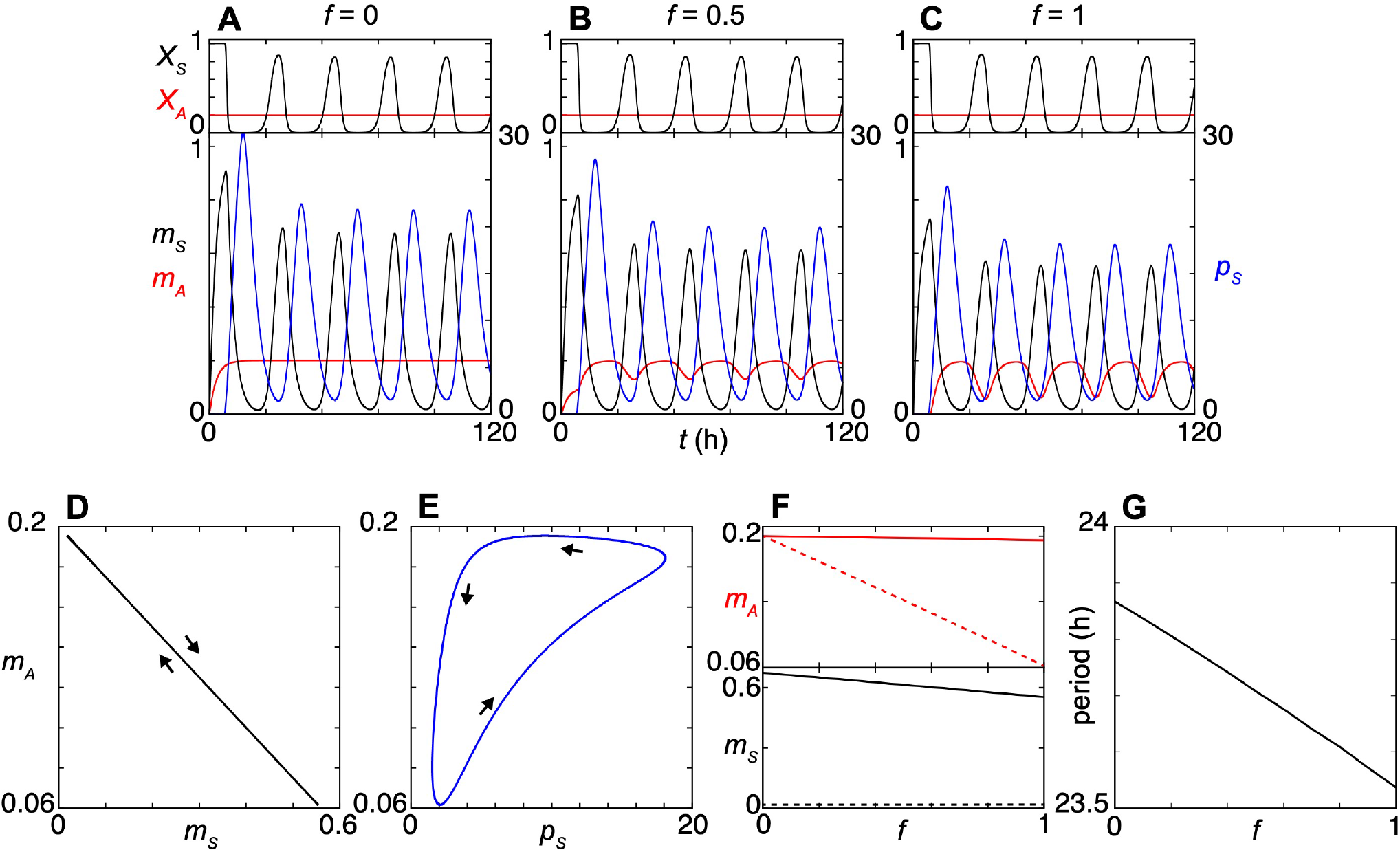
Characteristics of *Per2AS* and *Per2* expression patterns with our new transcriptional interference model. (A-C) Dynamics of *Per2* transcriptional activity (*X*_*S*_: black, top), *Per2AS* transcriptional activity (*X*_*A*_ : red, top), *Per2* mRNA (*m*_*S*_: black, bottom), *Per2AS* (*m*_*A*_: red, bottom), and PER2 protein (*p*_*S*_: blue, bottom) when the probability of RNAP collision and detachment *f* = *f*_*S*_ = *f*_*A*_ is 0 (A), 0.5 (B) or 1 (C). (D - E) Limit cycle projection onto (D) *m*_*S*_-*m*_*A*_ or (E) *p*_*S*_-*m*_*A*_ concentration spaces when *f* = 1. Arrows indicate the direction of trajectories. (F) Relationship between the probability of RNAP collision and detachment *f* and peak (solid) or trough (dotted) values of *m*_*A*_ (red: top), and *m*_*S*_ (black: bottom). (G) Period of *Per2* mRNA (*m*_*S*_) as a function of *f*.

### Our new model with transcriptional interference successfully describes the effect of Per2AS transcription perturbation

We next tested whether our new model would reproduce some of our experimental data. We first focused on the *Per2AS* mutant cell lines, in which a perturbation of the *Per2AS* promoter led to an increase of the *Per2AS* level while a decrease of the *Per2* level compared to a control cell (*Mosig et al., 2021*). In these mutants, the effect of *Per2AS* on *Per2* appears to be transcriptional, because the changes in the *Per2AS* level did not lead to the changes of the *Per2* level or circadian rhythms (*Mosig et al., 2021*).

To mathematically describe the changes observed in these mutants, we decreased the ratio of the rates between RNAP detachment and recruitment (i.e., *K*_*A*_ in Eq. (6b)) for *Per2AS* to represent the perturbation of the *Per2AS* promoter. Reduction of *K*_*A*_ increases the transcriptional activity of *Per2AS X*_*A*_ in Eq. (6b) and the production of *Per2AS* (the first term) in Eq. (8). In contrast, the reduction of *K*_*A*_ decreases the transcription of *Per2* mRNA (Eq. (7) first term). The effect of reducing *K*_*A*_ on *Per2* mRNA becomes stronger with the larger value of *f*_*S*_, which is the probability for RNAP collision and detachment on the *Per2* strand (Eq. (7)). Correspondingly, our numerical simulation demonstrated that the average *Per2* mRNA level becomes lower as *K*_*A*_ decreases and *f*_*S*_ = *f*_*A*_ = *f* increases (Fig. 3A). Our model also predicts that the period of *Per2* mRNA oscillation becomes shorter when *f* increases and *K*_*A*_ decreases (Fig. 3B). These results qualitatively corroborate with the experimental observations from *Per2AS* mutants (*Mosig et al., 2021*).

**Figure 3.**
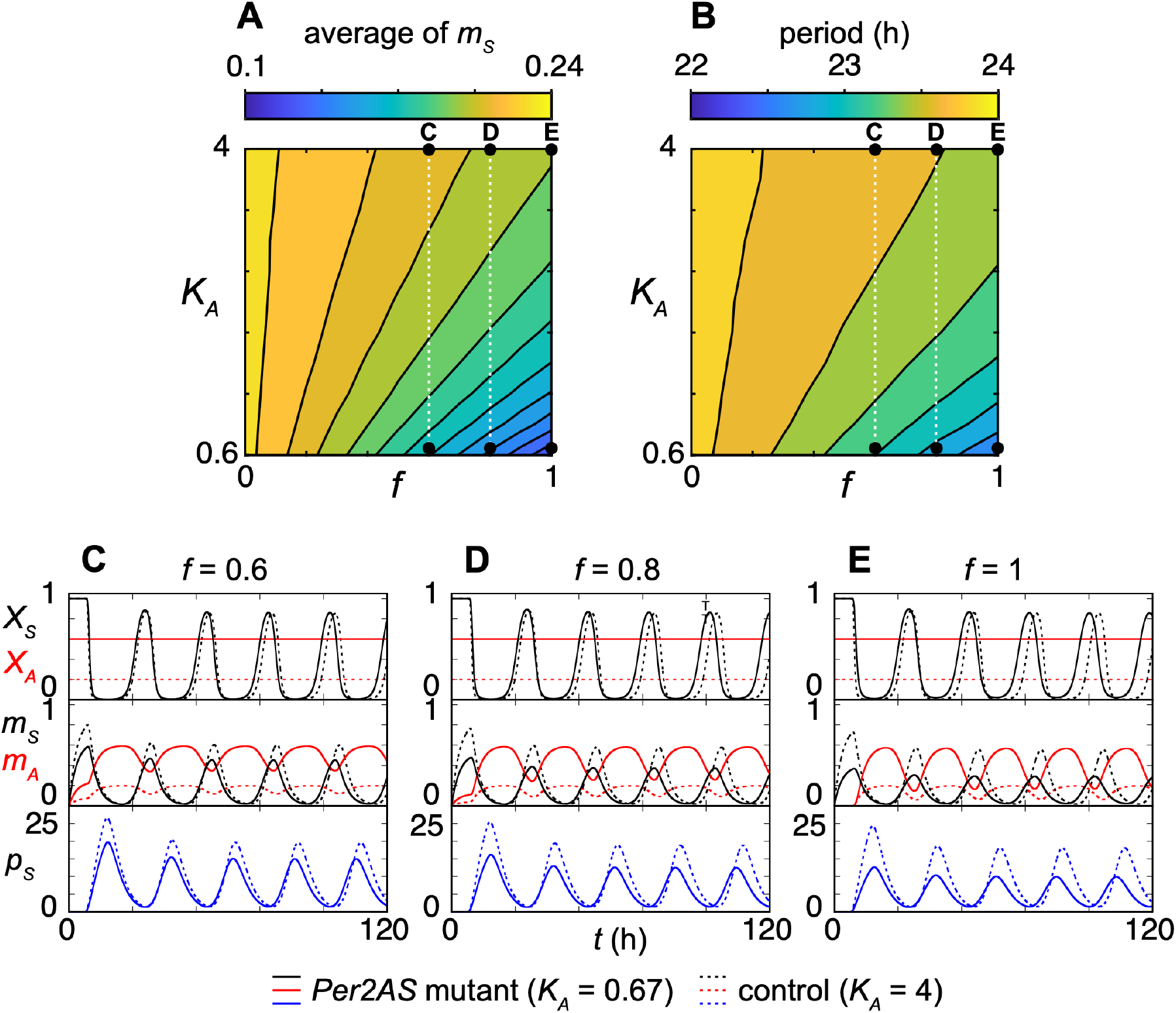
An increase in *Per2AS* transcription leads to the decrease of *Per2* and PER2 levels. (A-B) Contour plots of temporal averages of *Per2* mRNA (A) or period of *Per2* mRNA oscillation (B) showing their dependences on a probability of RNAP collision and detachment *f* and the ratio of the rates between RNAP dissociation and recruitment for *Per2AS* promoter *K*_*A*_. Black dots represent the parameter values of *f* and *K*_*A*_ used in (C-E). (C-E) Timeseries of transcriptional activity of *Per2AS* (*X*_*A*_), transcriptional activity of *Per2* (*X*_*S*_), the *Per2AS* transcript level (*m*_*A*_), *Per2* mRNA level (*m*_*S*_) and PER2 protein level (*p*_*S*_) with the parameter values described in Table 1. Broken lines indicate simulation with *K*_*A*_ = 4 representing wildtype. Solid lines indicate simulation with *K*_*A*_ = 0.67 representing *Per2AS* mutant. (C) *f* = 0.6, (D) *f* = 0.8, and (E) *f* = 1.

We next compared the dynamics of *Per2* mRNA and *Per2AS* between the mutant (*K*_*A*_ = 0.67) and the control (*K*_*A*_ = 4). When *K*_*A*_ decreases from 4 to 0.67, the level of *Per2AS* increased by 3-fold regardless of the value of *f* in the mutant (*m*_*A*_ in Fig. 3C-E), consistent with our experimental observations (*Mosig et al., 2021*). We observed a minimal impact of reduction of *K*_*A*_ on the amplitude of *X*_*S*_ (i.e., the transcriptional activity of *Per2*), while the peak level of *Per2* mRNA (Fig. 3C: *m*_*S*_) and PER2 (Fig. 3C: *p*_*S*_) were modestly decreased when *f* = 0.6. As we increase the probability of RNAP collision and detachment equally on both strands (i.e., *f* = 0.8 or 1.0), the amplitudes of *Per2* mRNA (*m*_*S*_) and PER2 protein (*p*_*S*_) in the mutant become smaller (Fig. 3D, E). In contrast, the amplitude of *Per2AS* increased with the increase in *f* in the mutant (Fig. 3C-E). Thus, our new model with transcriptional interference can describe an experimental observation, in which the perturbation of *Per2AS* promoter results in the increased of *Per2AS* and reduction of *Per2* mRNA.

### Our new model with transcriptional interference also successfully describes the effect of Per2 knock-down

One of the experimental observations that puzzled us was that a reduction of *Per2* mRNA levels by a gene knock-down led to a decreased level of *Per2AS* (*Mosig et al., 2021*). In our previous model (*Battogtokh et al., 2018*), we assumed that *Per2* and *Per2AS* mutually repress each other and the repression was mediated by their RNA or protein products. Under this condition, we expected that the *Per2AS* levels would be higher when *Per2* was knocked-down. However, this is in direct conflict to our experimental observations, prompting us to test whether the new mutual repression model with transcriptional interference can explain these experimental observations if the repression was independent of the *Per2* and *Per2AS* transcripts.

To mathematically describe the effect of *Per2* mRNA reduction by a gene knock-down, we first modified the second term of Eq. (7) from −*m*_*S*_(*t*) to −(1 + α)*m*_*S*_(*t*), where α represents the additional mRNA degradation caused by the shRNA-mediated gene knock-down (*Moore et al., 2010*). When we increased the degradation rate of *Per2* mRNA and shorten the half-life of *Per2* mRNA (*t*_*h*_ = (log 2)/*μ*_*S*_(1 + α)) from 2 h to 0.4 h, the average and amplitude of *Per2* mRNA level and those of PER2 protein level both decreased to less than half (Fig. 4A-C: *m*_*S*_ and *p*_*S*_). Decrease in PER2 protein level reduces the transcription of *Per2AS* in the first term of Eq. (8). Accordingly, as probability of RNAP collision and detachment *f* increased, the average *Per2AS* level in *Per2* knock-down condition became smaller than that in control condition in simulation (Fig. 4A-C). These results indicate that the mutual repression between *Per2AS* and *Per2* can still explain a counterintuitive experimental data, in which the knock-down of *Per2* led to the reduced level of *Per2AS* when the mutual repression is implemented as transcriptional interference.

**Figure 4.**
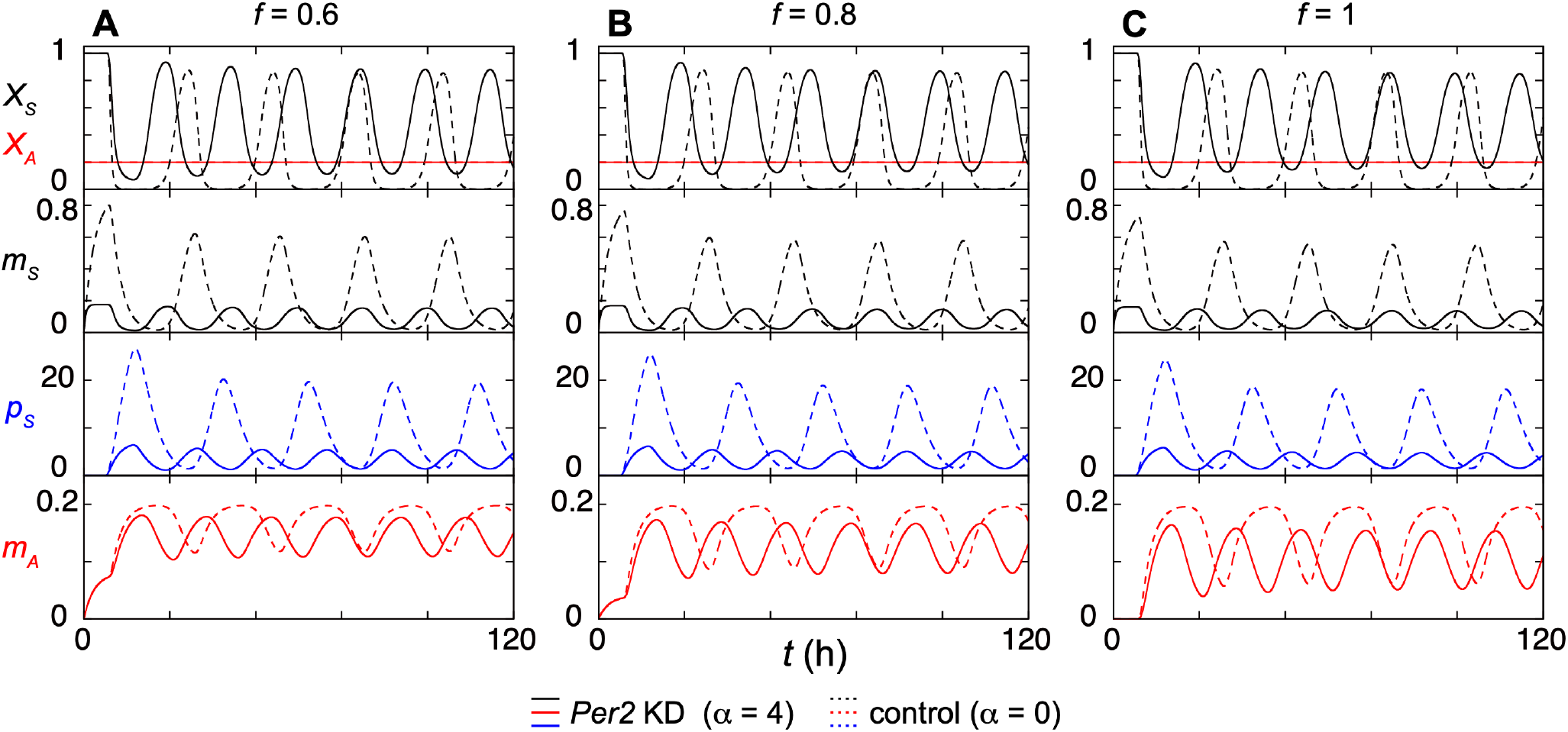
An increase in *Per2* degradation leads to a decrease of *Per2AS* transcript level. (A-C) Timeseries of the *Per2* transcriptional activity (*X*_*S*_), *Per2AS* transcriptional activity (*X*_*A*_), *Per2* mRNA level (*m*_*S*_), PER2 protein level (*p*_*S*_) and *Per2AS* transcript level (*m*_*A*_), when the probability of RNAP collision and detachment is *f* = 0.6 (A), 0.8 (B), or 1 (C). Dotted lines are simulation results for control (2 h of *Per2* mRNA half-life, *α* = 0) while solid lines are simulation results for *Per2* knock-down condition (0.4 h of *Per2* mRNA half-life, *α* = 4).

### Per2 overexpression does not decrease Per2AS

We next sought to test our new model with transcriptional interference further whether PER2 regulates *Per2AS* experimentally. When we overexpressed *Per2* exogenously using a plasmid, we found that the level of *Per2AS* was unaltered (Fig. 5A-B), which incosistent with the prediction from our model (Eq. (8)). In addition, the endogenous *Per2* transcript level was also unchanged (Fig. 5C). These data suggest that the exogenously expressed *Per2* mRNA and PER2 do not have any impact on *Per2AS*, or our current model is too simplified and additional factors are required to fully explain this experimental observation.

**Figure 5.**
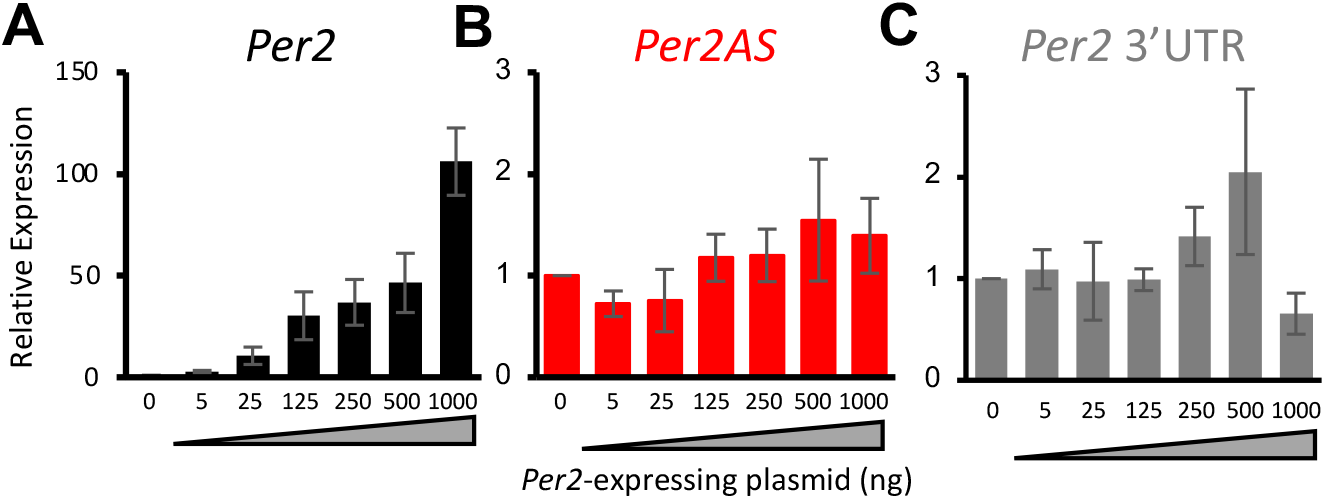
Overexpression of *Per2* does not alter the *Per2AS* level. (A-C) The levels of *Per2* (black; A) and *Per2AS* (red; B), and *Per2* 3’UTR (C) when PER2 was overexpression in AML12 cells. Data were normalized with the expression level of *36B4* and represent mean ± SEM (n=3-12) and the RNA levels of the control cells (0 ng of DNA transfection) were set to 1.

## Discussion

In this study, we developed a new mathematical model that mechanistically described the mutually repressive relationship between *Per2* and *Per2AS* via transcriptional interference. Our new model is mathematically viable and generates the rhythmicity of *Per2*, PER2, and *Per2AS*, with a period of 23.87 hour as well as the antiphasic oscillations between *Per2* and *Per2AS* (Fig. 2). The simulation results from the model are also consistent with the experimental observations including the counterintuitive ones (Figs. 3, 4). However, we also found new experimental observations, which is that the exogenous overexpression of *Per2* mRNA and PER2 protein have no impact on *Per2AS* (Fig. 5), cannot be explained by the current model with transcriptional interference. We think that the inconsistency between theory and experiment is presumably because the overexpression of *Per2* is independent from the endogenous system described in our model, because in our model PER2 protein represses endogenous *Per2* transcription (Eq. (7)) and thereby activates *Per2AS* through relieving transcriptional interference by *Per2* (Eq. (8)). It is also possible that PER2 alone is not sufficient to repress its own transcription and needs CRY1/2 to interact with and inhibit the activity of CLOCK-BMAL1 (*Griffin et al., 1999, Kume et al., 1999, Lee et al., 2001*). It is also plausible that *Per2AS* is regulated by other components besides *Per2*. In fact, we previously reported that the expression pattern of *Per2AS* is significantly dampened in *Bmal1*^*-/-*^ and *Cry1*^*-/-*^*Cry2*^*-/-*^ mice, while exhibits higher amplitude in *Nfil3*^*-/-*^, *Nd1d1*^*-/-*^*Nr1d2d*^*-/-*^, *Dbp*^*-/-*^*Tef*^*-/-*^*Hlf*^*-/-*^ mice (*Miao et al., 2022*), indicating that the expression of *Per2AS* is activated by *Bmal1* and/or *Cry1/2* and repressed by *Nfil3, Nr1d1/2*, or *Dbp/Tef/Hlf*. Their protein products may directly regulate the expression of *Per2AS* independent of PER2 by getting recruited to the promoter of *Per2AS* and or indirectly regulate *Per2AS* by regulating PER2 or other ‘core clock’ components.

To reconcile all these data, we propose a new topology that includes an additional feedback loop that connects PER2 and *Per2AS* via what we call Factor X (Fig. 6). Factor X would be either repressed by PER2 and activates *Per2AS* or activated by PER2 and represses *Per2AS* (Fig. 6), and this additional loop would serve as a buffer to the change the *Per2AS* level by transcriptional interference from *Per2*. If the former, the strongest candidates for Factor X would be either BMAL1 or CRY1/2, because the level of *Per2AS* was almost completely abolished in *Bmal1*^*-/-*^ and *Cry1*^*-/-*^*Cry2*^*-/-*^ mice (*Miao et al., 2022*). If the latter, the strongest candidates would be NFIL3, DBP, or REV-ERBα/β, given that they repress *Per2AS* (*Miao et al., 2022*). These possibilities can be best tested in an experimental setting and the outcomes can be further incorporated into our models to not only ultimately better understand the function of *Per2AS* but also how *Per2AS* contributes to the regulation of the robustness of the circadian clock system.

**Figure 6.**
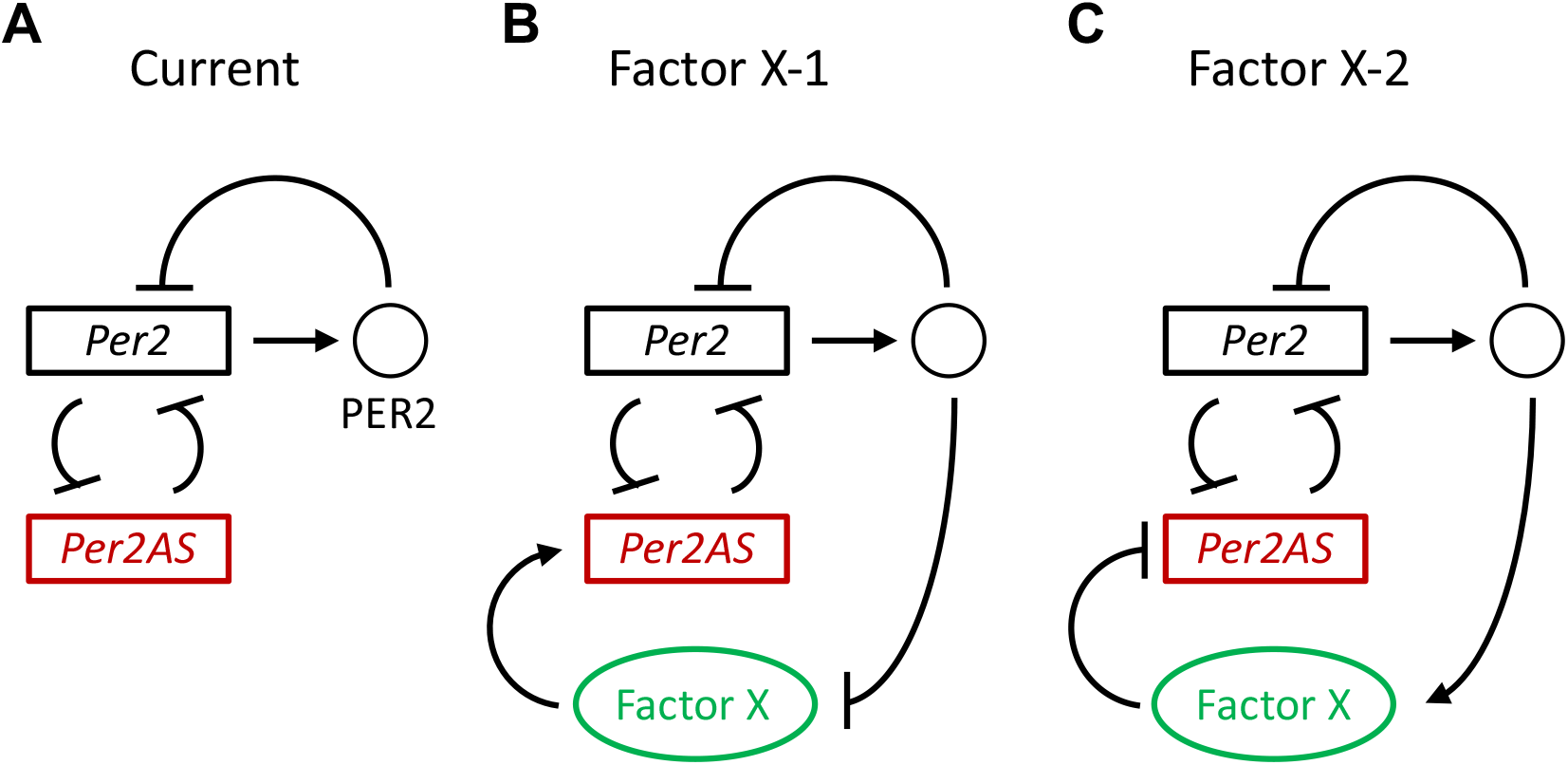
Proposed new topology. (A) Regulatory topology adopted in Eqs. (5), (7), (8). (B) Factor X activates *Per2AS* and it is repressed by *Per2*. If exogeneous PER2 protein represses *Per2*, transcriptional interference of *Per2AS* by *Per2* becomes weak. However, the PER2 protein also represses Factor X, activation of *Per2AS* by the Factor X also becomes weak. As a result, change in *Per2AS* level is buffered. (C) Factor X represses *Per2AS* and it is activated by *Per2*. In this case, exogeneous PER2 protein represses *Per2AS* via activating Factor X. Hence, activation of *Per2AS* by weakened transcriptional interference is balanced.

Overall, our new model supported the mutual repression via transcriptional interference between *Per2AS* and *Per2*, while it also suggested that it needs to be further refined to fully describe the intricate genetic network of the mammalian circadian clock system involving *Per2AS*. Because of the simplicity of our model, it can also be extended to understand the impact of mutual repression between any antisense-sense pairs (both mRNAs and non-coding RNAs) that use transcription interference as a regulatory mechanism. Mathematical models that describe transcriptional interference are currently scarce, and our models can serve as a foundation to understand the function of mutual repression in any genetic circuit, including both synthetic and natural, even outside of the circadian rhythms.

## Materials and Methods

### Derivation of equations for dimensionless variables Eqs. (3)-(5)

Here we derive equations for dimensionless variables Eqs. (3)-(5) from the corresponding dimensional equations:

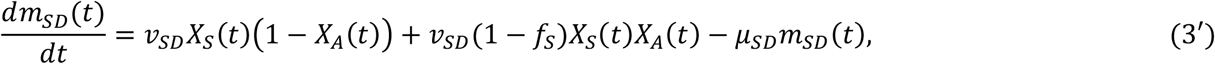

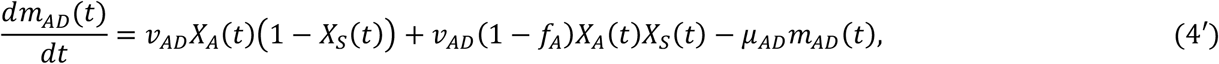

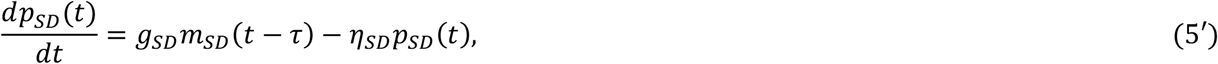

where subscripts *D* in the variables and parameters denote that they have a unit. Note that transcriptional activities *X*_*S*_ and *X*_*A*_ are the numbers without units. We define dimensionless variables *m*_*S*_, *m*_*A*_ and *p*_*S*_ by using a concentration scale *v*_*SD*_/*μ*_*SD*_ (i.e., transcription rate of *Per2* mRNA divided by its degradation rate) as:

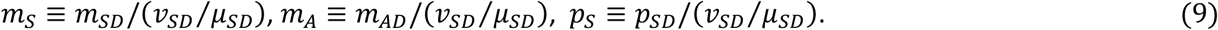

By substituting Eq. (9) into Eqs. (3’)-(5’), we obtained Eqs. (3)-(5) with dimensionless parameters:

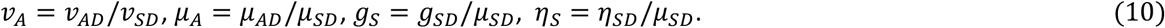

We keep a timescale of the system 1/*μ*_*SD*_ in the left-hand side of Eqs. (3)-(5). For notational simplicity, we omit subscript *D* of *μ*_*SD*_ and write *μ*_*SD*_ = *μ*_*S*_ in Eqs. (3)-(5).

### Numerical simulations

The delay differential equations, Eqs. (5), (7) and (8) were solved numerically with the dde23 function in MATLAB 2022b and 2023a (MathWorks) (see Data Availability Statement). Initial condition and history of the DDEs were *m*_*A*_(0) = *p*_*S*_(0) = 0, and *m*_*S*_(*t*) = 0 for −*τ* ≤ *t* ≤ 0. For the calculation of period, Eqs. (5), (7) and (8) were solved for 720 hours and last ∼10 time intervals between two successive peaks of *Per2* mRNA were averaged. Parameter values are listed in Table 1.

### Cell culture and DNA transfection

AML12 cells were grown in Dulbecco’s modified Eagle medium/F12 (Gibco, #11320033) with 10% FBS and 1% insulin–transferrin–selenium supplement (Gibco, #41400045) at 37°C with 5% CO2. A Per2 plasmid was transfected with polyethylenimine (Polysciences, #233966-100) using Opti-MEM (Gibco, # 31985070). AML12 cells and the *Per2* plasmid is a generous gift from Dr. Carla B. Green (UT Southwestern Medical Center).

### RT-qPCR

Total RNA was extracted with TRIZOL reagent (Life Technologies, #15596018) according to the manufacturer’s instructions and treated with TURBO DNaseI (Life Technologies, #AM2239). RNAs were then subjected to reverse transcription using SuperScript II (Life Technologies, # 18064014) or high-capacity cDNA reverse transcription kits (Applied Biosystems, #4368813). For *Per2AS* transcripts, cDNA was synthesized using strand-specific primer (5’-AGCTGGTCCAATGTCAGGAGG-3’). qPCR was performed using QuantiStudio 6 (Life Technologies) with SYBR Power Green (Applied Biosystems, #4367659). The primer sequences used in this study are as follows: m36B4-QF: 5’-cactggtctaggacccgagaag-3’, m36B4-QR: 5’-ggtgcctctgaagattttcg-3’, mPer2AS-QF: 5’-AGTAGAAAGAGGTAGGGAGGC-3’, mPer2AS-QR: 5’-TCATCTAAGGGTCTGGGAGAG-3’, mPer2_F: 5’-TGTGCGATGATGATTCGTGA-3’, mPer2_R: 5’-GGTGAAGGTACGTTTGGTTTGC-3’, mPer2_exon5_F: 5’-TATCGTGAAGAACGCGGATA-3’, mPer2_exon6_R: 5’-AGTGAAAGATGGAGGCCACT-3’.

## Data Availability Statement

MATLAB simulation codes are available from: https://github.com/uriukoichi/Per2AS_MATLABcodes.git

## Author contributions

KU – methodology, formal analysis, investigation, writing-original draft, writing – review & editing, visualization, project administration

JPHS – investigation, writing – review & editing, visualization,

SK – conceptualization, investigation, writing-original draft, writing – review & editing, supervision, project administration, funding acquisition

## Acknowledgement

The authors thank Dr John Tyson for the critical reading of the manuscript. The authors also thank Evan Littleton and Lin Miao for invaluable discussion.

